# Sulfation of glycosaminoglycans depends on catalytic activity of a lithium-inhibited phosphatase

**DOI:** 10.1101/2021.06.24.449779

**Authors:** Brynna S. Eisele, Zigmund A. Luka, Alice J. Wu, Fei Yang, Andrew T. Hale, John D. York

## Abstract

Golgi-resident bisphosphate nucleotidase 2 (BPNT2) is a member of a family of magnesium-dependent/lithium-inhibited phosphatases that share a three-dimensional structural motif that directly coordinates metal binding to effect phosphate hydrolysis. BPNT2 is responsible for the breakdown of 3’-phosphoadenosine-5’-phosphate (PAP), a by-product of glycosaminoglycan (GAG) sulfation. Disruption of BPNT2 in mice leads to skeletal abnormalities due to impaired GAG sulfation, especially chondroitin-4-sulfation. Mutations in *BPNT2* have also been found to underlie a chondrodysplastic disorder in humans. The precise mechanism by which loss of BPNT2 impairs sulfation remains unclear. Here, we utilize an *in vitro* approach using mouse embryonic fibroblasts (MEFs) to test the hypothesis that catalytic activity of BPNT2 is required for GAG sulfation. We show that a catalytic-dead *Bpnt2* construct (D108A) does not rescue impairments in intracellular or secreted sulfated GAG, including decreased chondroitin-4-sulfate, present in *Bpnt2*-knockout MEFs. We also demonstrate that missense mutations in *Bpnt2* which are adjacent to the catalytic site (and known to cause chondrodysplasia in humans) recapitulate defects in overall GAG sulfation and chondroitin-4-sulfation in MEF cultures. We further show that treatment of MEFs with lithium inhibits GAG sulfation, and that this effect depends on the presence of BPNT2. This work demonstrates that the catalytic activity of an enzyme potently inhibited by lithium can modulate GAG sulfation and therefore extracellular matrix composition, revealing new insights into lithium pharmacology and the pathophysiology of psychiatric disorders responsive to lithium.

## Introduction

Sulfation is a ubiquitous biological process in eukaryotes wherein a sulfate group from phosphoadenosine-phosphosulfate (PAPS, the universal sulfate donor) is transferred to a target substrate by sulfotransferase enzymes. This reaction yields the by-product 3’-phosphoadenosine-5’-phosphate (PAP), which is further catabolized to 5’-adenosine monophosphate (AMP) by the bisphosphate nucleotidases (BPNT1 and BPNT2)^1–5^. BPNT1 is localized to the cytoplasm, where sulfation of small molecules (such as hormones and xenobiotics) occurs^2^. BPNT2 (previously known as LPM, IMPAD1, gPAPP) is localized to the Golgi, which is the site of glycosaminoglycan (GAG) sulfation^1^. Sulfated GAGs are important components of the extracellular matrix that serve important structural roles and facilitate cell-to-cell signaling^6^.

A role for BPNT2 in regulating GAG sulfation was discovered through the generation of *Bpnt2*-knockout mice. These mice die in the perinatal period, but pups have shortened limbs indicative of chondrodysplasia^1,7^. Further analysis of tissue from *Bpnt2*-knockout pups on embryonic day 18.5 (E18.5) identified a significant decrease in GAG sulfation, particularly of chondroitin-4-sulfate^1^, which is necessary for the production of the cartilage matrix that precedes endochondral ossification of long bones. Subsequent studies by other groups identified an autosomal recessive human disorder caused by mutations in *BPNT2* and characterized by chondrodysplasia^8–10^, reminiscent of the knockout mouse phenotype, as well as other disorders of GAG sulfation^11^. However, the mechanism by which loss or mutation of *Bpnt2* impairs GAG sulfation is not currently known.

The BPNT enzymes are members of a family of magnesium-dependent, lithium-inhibited phosphatases^1,3,12^. The precise mechanism and location of lithium-mediated inhibition of these enzymes was recently reported, establishing this family of enzymes as direct targets of lithium^13^. Lithium has been used for more than a half-century as a treatment for psychiatric disorders^14^, but its therapeutic mechanism remains unclear^15^. Prior work suggests that lithium may modulate chondroitin sulfate^16,17^, but the mechanisms by which this could occur remain unknown. BPNT2 is a known modulator of chondroitin sulfation and a direct target of lithium^1^. Thus, BPNT2 inhibition is a candidate mechanism for lithium’s purported effects on chondroitin sulfate, and this inhibition may contribute to the therapeutic consequences or side effects of lithium treatment. However, previous studies have not established whether the consequences of *Bpnt2*-knockout originate from a loss of BPNT2’s catalytic activity (namely, the conversion of PAP to 5’-AMP) or from another non-catalytic function.

The objective of this study was to investigate whether the loss of BPNT2’s catalytic function underlies the chondrodysplastic phenotype observed in mice and humans. We established a model system to study BPNT2’s effects on GAG sulfation *in vitro* using embryonic fibroblasts cultured from *Bpnt2*-knockout mice. We then utilized genetic complementation to examine the impact of mutations in *Bpnt2* on GAG sulfation. We studied 3 *Bpnt2* mutations, which happen to be in close proximity to the active site/metal-binding domain (1 which ablates catalytic activity, 2 which are known to cause chondrodysplasia in huamns^8^). We demonstrate herein that these mutations impair GAG sulfation. We further show that treatment of MEF cultures with lithium chloride decreases GAG sulfation; these effects are dependent on the presence of BPNT2, consistent with BPNT2 being an *in vivo* target of the drug.

## Results

### Generation of an in vitro model system to analyze GAG sulfation

BPNT2 is a Golgi-resident protein which has a demonstrated role in Golgi-localized sulfation reactions. The major components of this sulfation pathway are illustrated in **Figure 1A**. Loss of BPNT2 is known to impair the upstream sulfation of GAGs, but previous studies have not established whether this effect stems from the loss of BPNT2 catalytic activity or another non-catalytic function. To investigate this mechanism, we first sought to generate an immortalized cell system to study the function of BPNT2 with respect to alterations in GAG sulfation. We elected to use embryonic fibroblasts because: 1. These cells were easily attainable from the *Bpnt2-*knockout mouse line developed by our laboratory (Jackson #012922); 2. MEFs derive from mesenchyme, which give rise to connective tissues which are primarily responsible for GAG synthesis *in vivo*, and 3. MEFs can be readily immortalized to facilitate genetic manipulations and prolonged study. MEFs were harvested on embryonic day 12.5 of pregnancies resulting from *Bpnt2*-heterozygous crosses. *Bpnt2* wild-type (WT) and knockout (KO) MEFs obtained from littermates were then immortalized by lentiviral expression of SV40 T antigens. Absence of BPNT2 in these lines was further confirmed by analyzing mRNA expression using quantitative PCR, and by immunoblotting for BPNT2 protein (**Figure 1B**). For our analyses, the most relevant property of the cells was their ability to synthesize and secrete sulfated GAG, the most abundant of which is chondroitin sulfate. The cells primarily responsible for chondroitin sulfate production *in vivo* are chondrocytes, but chondrocytes are not well suited to long-term culture involving repeated passaging. However, chondrocytic properties can be induced in cells of mesenchymal origin by maximizing cell-cell contact in cultures and supplementing media with certain growth factors^18^. This method of culturing immortalized MEFs allowed us to investigate the effects of BPNT2 on GAG sulfation *in vitro*, without having to repeatedly harvest primary cells, and allowed us to manipulate gene expression to generate stable cell lines expressing mutant versions of BPNT2. *Bpnt2*-KO MEFs display decreased total GAG sulfation as measured by dimethylmethylene blue (DMMB) assay assay (**Fig 1C, left panel**), normalized to cell number. KO MEFs also secrete fewer sulfated GAGs into the culture medium (**Fig 1C, right panel**). We were also interested in whether these cells displayed measurable alterations in PAP level, which might be expected in the absence of BPNT2, but we did not detect any difference in PAP between WT and KO cells. Nonetheless, the alterations in sulfation in immortalized MEFs recapitulate impairments in sulfation seem in *Bpnt2*-KO mice. We therefore deemed this an appropriate model for our investigations.

**Figure 1.**
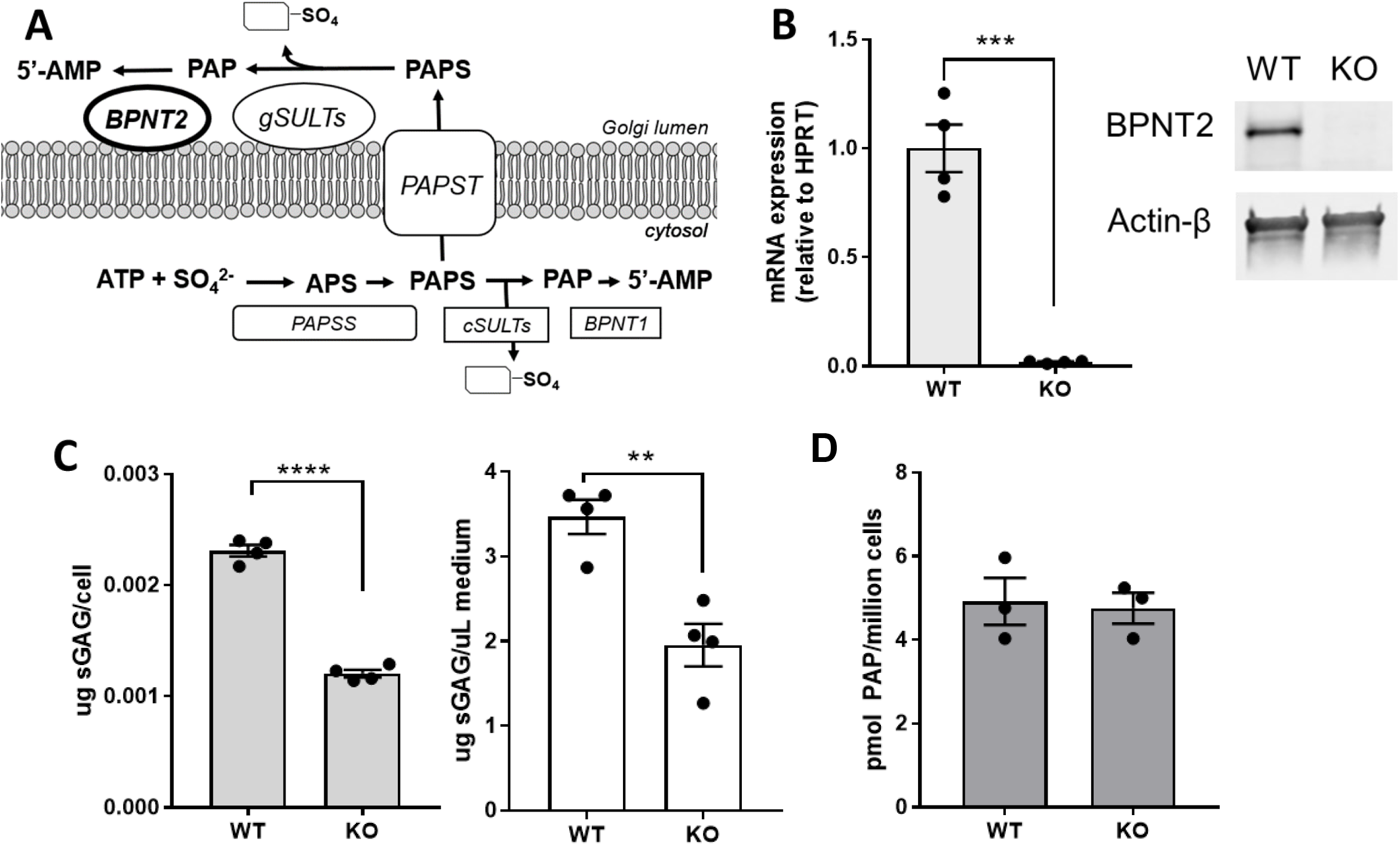
Loss of *Bpnt2* impairs glycosaminoglycan sulfation, but does not alter PAP level. A. Illustration of intracellular sulfation pathways, wherein BPNT2 hydrolyzes PAP, a by-product of sulfation, to AMP. B. Absent expression of *Bpnt2* mRNA (\*\*\**p*=*0.0001*) and protein in immortalized *Bpnt2*-KO MEFs. C. *Bpnt2*-KO MEFs exhibit decreased levels of both intracellular (left, \*\*\*\**p*<*0.0001*) and secreted (right, \*\**p*=*0.0034*) sulfated glycosaminoglycans, as determined by DMMB assay. D. *Bpnt2*-KO MEFs do not show changes in PAP level relative to WT cells. Bars show mean +/− SEM. Significance analyses are results of unpaired student’s *t* test (2-sided). **ATP:** adenosine triphosphate, **APS:** adenosine phosphosulfate, **PAPSS:** PAPS synthase, **PAPS:** 3’,5’-phosphoadenosine phosphosulfate, **PAP**: 3’,5’-phosphoadenosine phosphate, **AMP**: 5’-adenosine monophosphate, **cSULTs**: cytosolic sulfotransferases, **PAPST**: PAPS transporter, **gSULTs**: Golgi-resident sulfotransferases.

### Bpnt2 mutations that cause chondrodysplasia are located near the metal-binding/catalytic domain

Murine BPNT2’s three-dimensional core structural motif (which defines the family of lithium-inhibited phosphatases) has been simplified and represented graphically in **Figure 2A**. This region of BPNT2 is highly conserved across species^8^. Three aspartic acid (D) residues provide a negatively-charged environment conducive to the binding of positively-charged metal cations: divalent magnesium is a necessary cofactor for phosphate hydrolysis, whereas monovalent lithium inhibits this hydrolysis^13^. Mutation of the first aspartic acid which composes this pocket (D110^human^/D108^mouse^) to alanine renders family members catalytically inactive^13,19^. Interestingly, two missense mutations in *Bpnt2* localized near this locus are known to cause chondrodysplasia in humans^8,9^. A summary of these mutations is shown in **Figure 2B**. These mutant versions of *Bpnt2* have previously been predicted to have effects on the enzymatic activity, due to localization near the presumed active site, based on structural comparisons to BPNT1 (PDB: 2WEF)^8^, as no structure has yet been determined for BPNT2. Indeed, one of these mutations, D177N^human^/D175N^mouse^, is in another of the three aspartic acid residues that compose the negatively-charged pocket, while T183P^human^/T181P^mouse^ is located just 6 amino acids further downstream.

**Figure 2.**
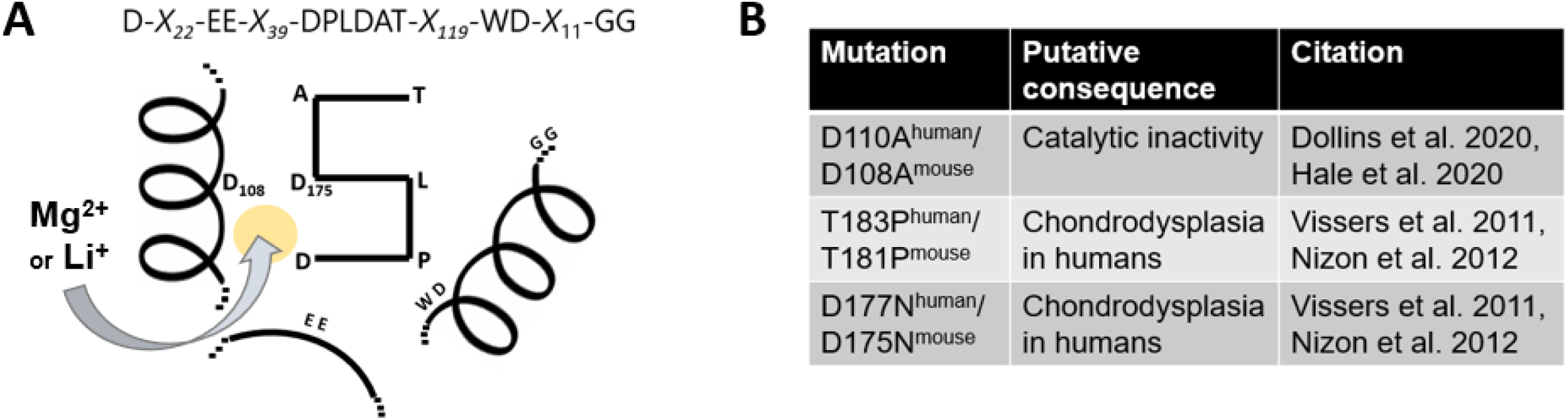
Summary of *Bpnt2* mutations in relation to the metal-binding/catalytic domain. A. Graphical simplification of metal-binding/catalytic domain defining magnesium-dependent/lithium-inhibited phosphatases; numbered amino acids are those in murine *Bpnt2*; yellow circle represent metal-binding pocket, where magnesium binds under active conditions and lithium binds under inhibitory conditions. B. Table showing mutations investigated herein, which are in close proximity to the metal-binding structural motif.

### Expression of Bpnt2 mutants revealed a novel N-glycosylation locus on chondrodysplasia-associated mutant Bpnt2^D175N^

We next utilized site-directed mutagenesis and a retroviral expression system to generate MEF lines which exclusively express mutant forms of murine *Bpnt2*: *Bpnt2*^D108A^, *Bpnt2*^T181P^, and *Bpnt2*^D175N^. We also generated a wild-type *Bpnt2*-complemented cell line (KO+WT) as a control, as well as WT and KO lines transduced with empty vector (EV) control retrovirus. Success of viral transduction was determined by western blotting for BPNT2 (**Figure 3A**). Surprisingly, BPNT2^D175N^ mutant protein appeared to be partially shifted upward, displaying a second band of higher molecular weight relative to other BPNT2 isoforms. Native BPNT2 is localized to the Golgi, and like other membrane-associated proteins, it contains an N-glycosylation consensus sequence (N-X-S/T^20^) at N259^human^/N257^mouse^. The novel asparagine in BPNT2^D175N^ made us inquire as to whether an additional N-glycosylation locus was generated by this mutation, as a secondary glycosylation event could explain the increased molecular weight of the detected protein. **Figure 3B** shows a section of cDNA for both human and mouse *Bpnt2*. In both human and mouse, the mutation of this aspartic acid to asparagine results in the generation of an N-glycosylation consensus sequence: N-A-T. To test whether the heavier band was indeed due to an additional N-glycosyl modification, protein extracts from cell lines were treated with PNGaseF. Native BPNT2’s one glycosylation site at N259^human^/N257^mouse^ is cleaved with PNGaseF treatment, resulting in a downward shift of the protein. Treatment of BPNT2^D175N^ with PNGaseF cleaves both N-glycosyl groups, eliminating the double band and producing a protein of the same size as those seen in all other PNGaseF-treated cell lines (**Figure 3C**). The generation of this additional N-glycosylation site has not been previously described in association with this mutation.

**Figure 3.**
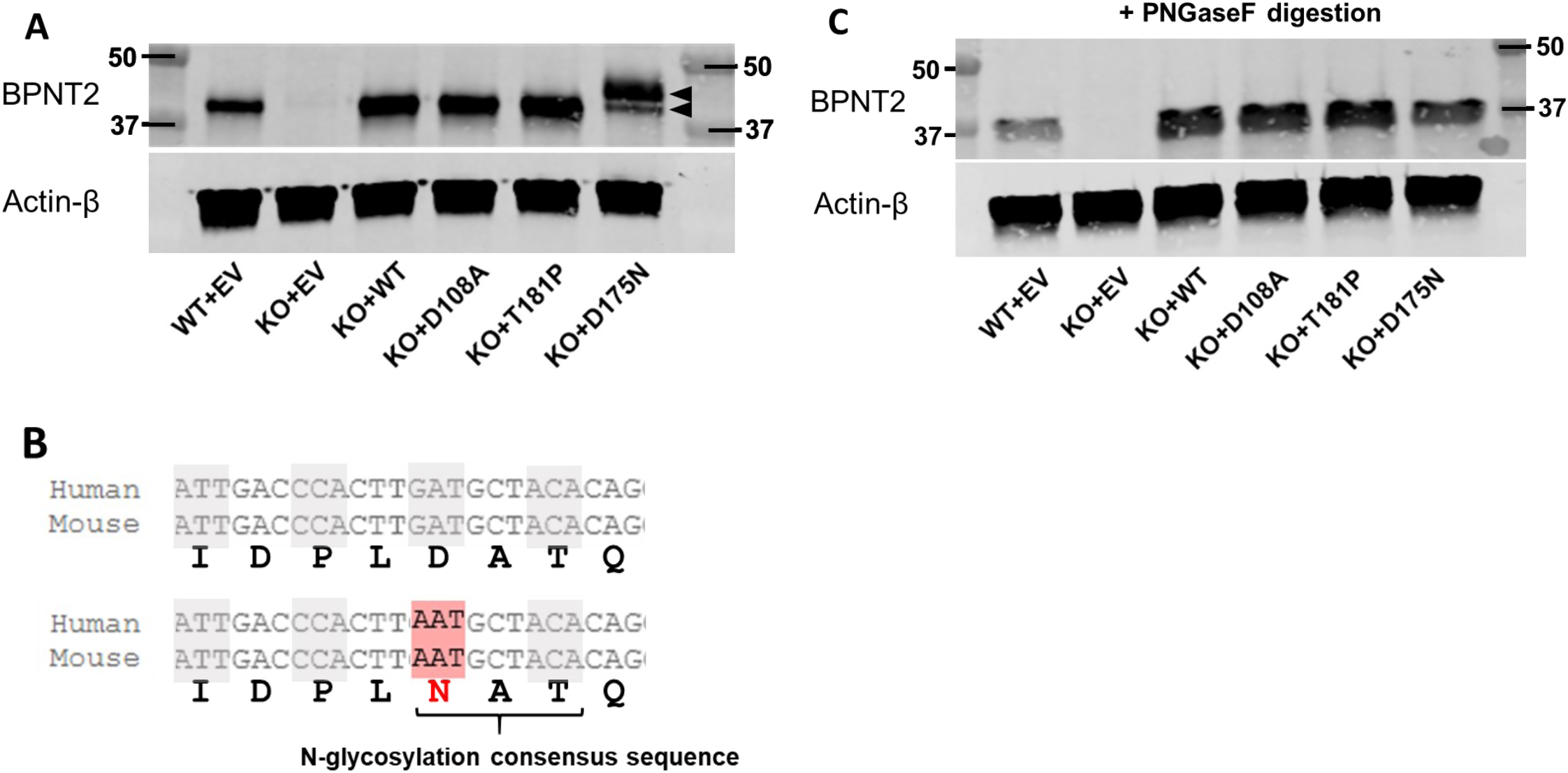
Generation of mutant *Bpnt2* MEF lines. A. Blot for BPNT2 and actin on protein extracted from *Bpnt2* MEF lines; arrows in D175N lane denote 2 bands, representing singly and doubly glycosylated protein. B. A selection of *Bpnt2* sequence from mouse and human *Bpnt2*. In both, the D175N/D177N mutation results in generation of an N-glycosyl consensus sequence. C. Blot of proteins extracted from MEF lines, treated with PNGaseF to remove N-glycosyl groups; PNGaseF removes both glycosyl groups, resulting in a single band for BPNT2-D175N. Note that all bands shift downward with PNGaseF, as native murine BPNT2 has one N-glycosylation site at N257.

### Wild-type Bpnt2 rescues impairments in overall sulfated GAGs, while mutant Bpnt2 constructs do not

We next sought to examine the consequences of these mutations on GAG sulfation. To facilitate the synthesis of GAGs and extracellular matrix components, we cultured three-dimensional cell pellets for each MEF line over a period of 7-14 days. Media was collected from pellets immediately prior to harvest. Total sulfated GAG from cell pellets was quantified using a colorimetric dimethylmethylene blue (DMMB) assay, then normalized to cell count. These results are depicted in **Figure 4A**. We observed a significant decrease in sulfated GAG in the knockout line, which was rescued by expressing wild-type *Bpnt2*. In contrast, expression of *Bpnt2*^D108A^ did not rescue this decrease, nor did *Bpnt2*^T181P^ or *Bpnt2*^D175N^. Level of secreted sulfated GAG in media (collected at pellet harvest) was also measured, and are shown in **Figure 4B**. Again, we observed a significant decrease in secreted/extracellular sulfated GAG in the knockout line, which was not rescued with expression of *Bpnt2*^D108A^, but did appear to be rescued with expression of other *Bpnt2* mutants.

**Figure 4.**
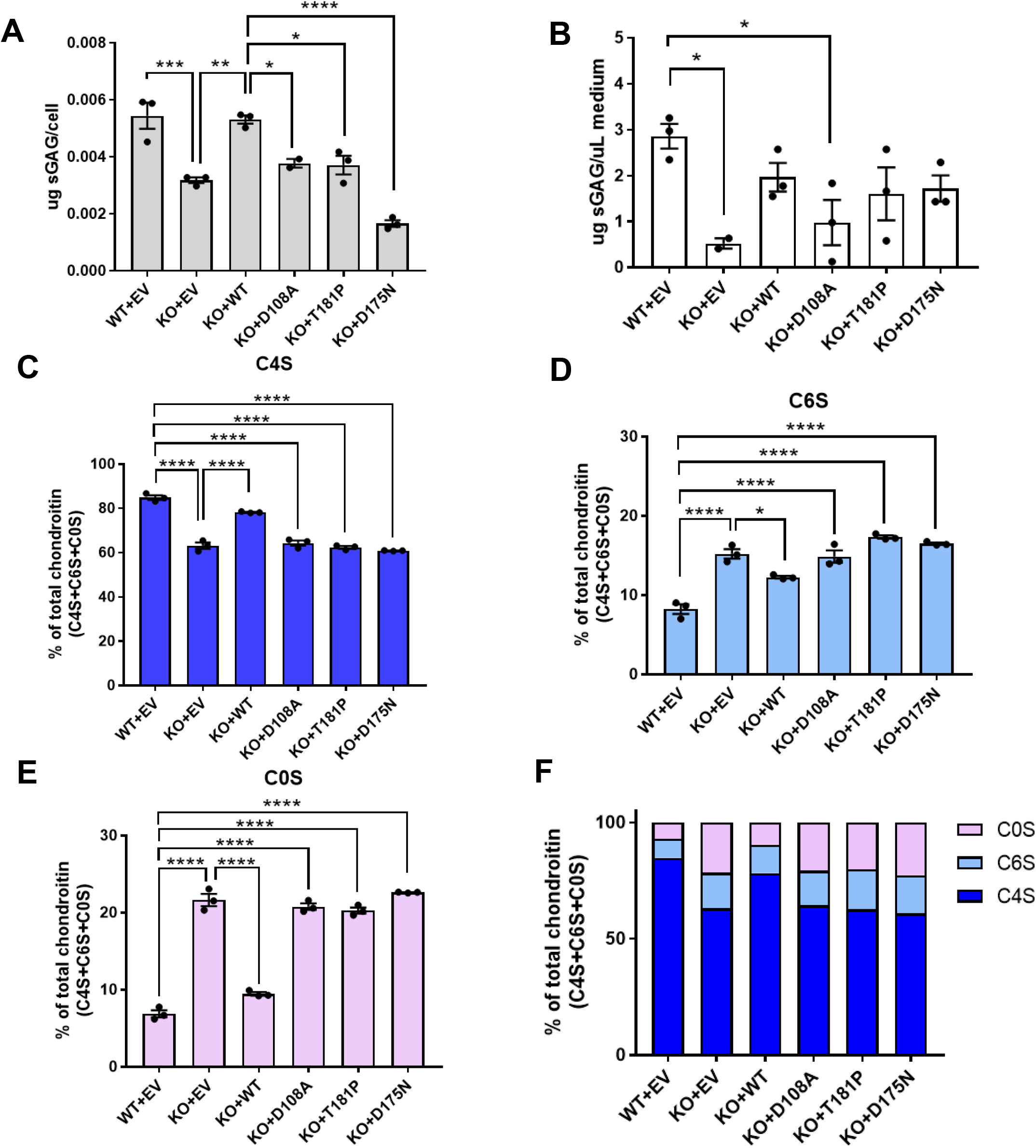
MEFs expressing mutated *Bpnt2* exhibit decreased sulfated glycosaminoglycans, including decreased chondroitin-4-sulfate. A. Sulfated GAG levels in *Bpnt2*-mutant cells, as measured by DMMB assay. B. Sulfated GAG levels in medium of *Bpnt2-*mutant cell cultures, as measured by DMMB assay. Alterations in C. chondroitin-4-sulfate, D. chondroitin-6-sulfate, and E. unsulfated chondroitin levels in *Bpnt2-* mutant cell cultures, as measured by chondroitin disaccharide HPLC. F. Summary of alterations in chondroitin-sulfation profile across *Bpnt2-*mutant lines. Error bars show mean +/− SEM. Denoted significance indicates results of Tukey’s post-hoc tests after significant one-way ANOVA. *p<0.05, **p<0.01, ***p<0.001, ****p<0.0001.

### Mutant Bpnt2 constructs do not rescue specific alterations in chondroitin sulfation

We next used high-performance liquid chromatography (HPLC) to evaluate specific alterations in chondroitin sulfation. Isolated glycosaminoglycans can be digested by chondroitinase enzymes to yield individual sulfated chondroitin disaccharides. These disaccharides can then be fluorescently labeled and resolved by HPLC to identify specific sulfation moieties. Previous work has demonstrated that *Bpnt2*-KO mouse embryos exhibit impairments in overall levels of 4-sulfated disaccharide (μdi-4S, or 4S) that correspond with increased levels of unsulfated chondroitin (μdi-0S, or 0S)^1^. We observed a decrease in the ratio of 4S to total chondroitin disaccharides (4S+6S+0S) in KO+EV MEFs relative to WT+EV MEFs, which was in large part rescued by complementing WT *Bpnt2* back into the line (KO+WT). However, this decrease was not rescued by expression of catalytic-dead *Bpnt2* or either chondrodysplasia-associated mutant *Bpnt2* (**Figure 4C**). Interestingly, we observed a correspondent increase in the ratio of chondroitin-6-sulfate (μdi-6S, or 6S) in KO cells, which was partially restored with WT complementation, and which remained elevated in mutant *Bpnt2* lines (**Figure 4D**). The increase in 6S has not been previously reported in association with *Bpnt2*-KO. However, overall ratios of unsulfated chondroitin are increased in KO cells, restored to near-WT levels with back-complementation, and significantly elevated in mutant *Bpnt2* lines (**Figure 4E**). These changes are summarized in **Figure 4F**.

### Lithium decreases intracellular and extracellular sulfated GAG, including chondroitin-4-sulfate

The discovery that the loss of BPNT2 catalytic function underlies impairments in overall GAG sulfation is particularly relevant because BPNT2 is potently inhibited by the psychopharmacologic agent lithium^1^. To test whether lithium impairs total GAG sulfation, we treated both WT and *Bpnt2*-KO MEFs with 10 mM lithium chloride (LiCl), using an equal concentration of sodium chloride (NaCl) as a control. We observed a decrease in intracellular GAG sulfation in lithium-treated WT MEFs, as compared to sodium-treated WT MEFs (**Figure 5A**). Loss of *Bpnt2* in cells treated with sodium resulted in a marked reduction in GAG sulfation similar to WT MEFs treated with lithium (**Figure 5A**). Importantly, treatment of *Bpnt2*-KO MEFs with lithium did not further reduce sulfation of GAGs (**Figure 5A**). Likewise, we observed a significant reduction in sulfation of GAGs secreted into the culture medium in lithium-treated as compared to control sodium-treated samples (**Figure 5B**). Again, *Bpnt2-*KO MEFs treated with LiCl did not exhibit any additional decrease in secreted sulfated GAG.

**Figure 5.**
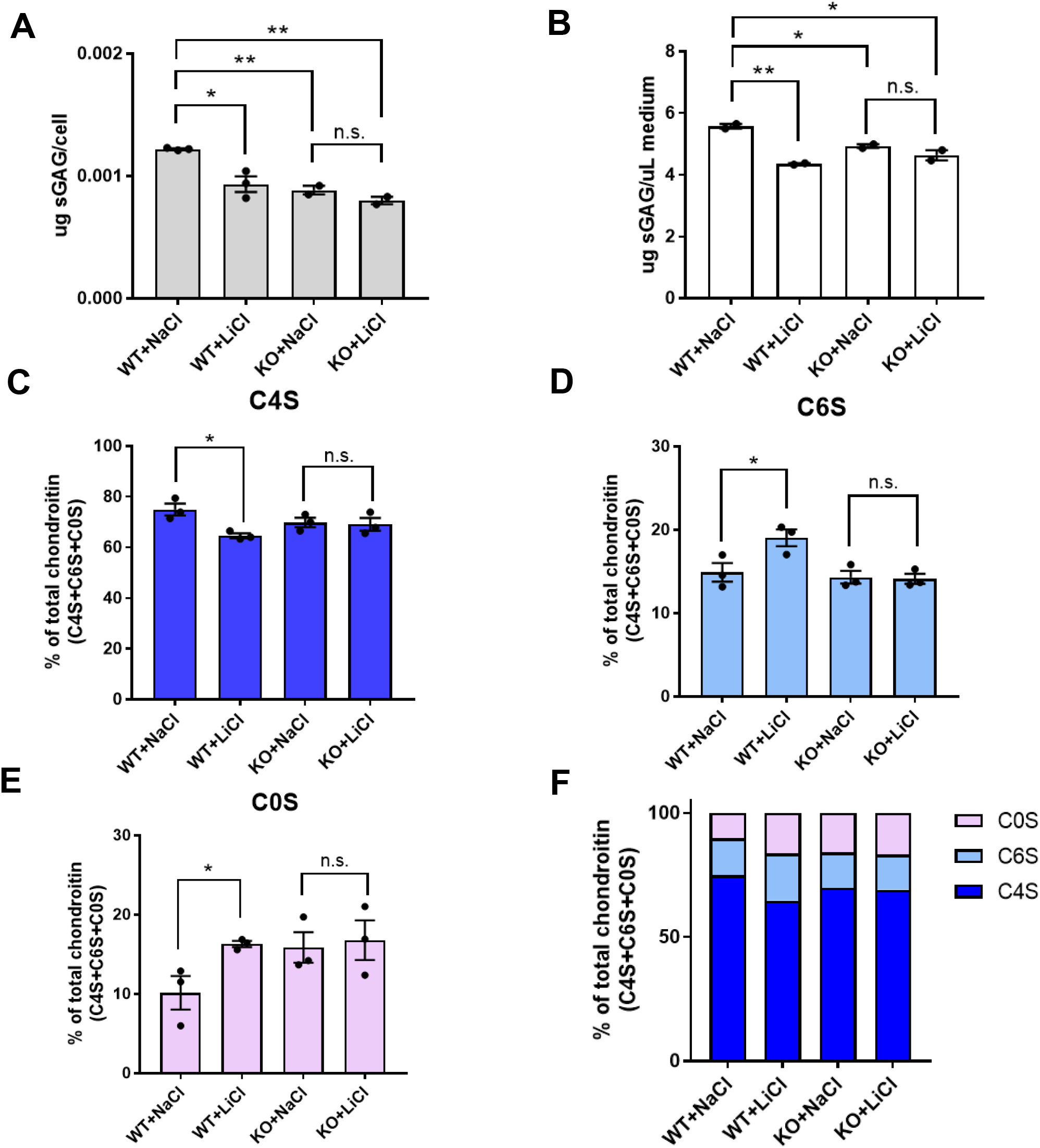
Lithium treatment decreases overall GAG sulfation, including chondroitin-4-sulfation, in wild-type cells, but not *Bpnt2-* knockout cells. A. Sulfated GAG analyses performed on cells treated with 10 mM NaCl or LiCl. B. Sulfated GAG analysis performed on culture medium of cells treated with 10mM NaCl or LiCl. Alterations in C. chondroitin-4-sulfate, D. chondroitin-6-sulfate, and E. unsulfated chondroitin levels in treatment groups, as measured by chondroitin disaccharide HPLC. F. Summary of alterations in chondroitin-sulfation profile across treatment groups. Error bars show mean +/− SEM. Denoted significance indicates results of two-sided student’s t-test. *p<0.05, **p<0.01, n.s. = not significant.

We next used HPLC to resolve chondroitin disaccharides in WT and *Bpnt2-*KO cells treated with lithium, and we observed a significant decrease in 4S (**Figure 5C**) in LiCl-treated WT cells alongside significant increases in 6S (**Figure 5D**), and 0S (**Figure 5E**). *Bpnt2*-KO cells did not exhibit these alterations when treated with LiCl, demonstrating that lithium does not have any additional effects on chondroitin sulfation patterns in cells that lack *Bpnt2*. Collectively, the similarity between sulfation patterns in LiCl-treated cells and *Bpnt2*-KO cells can be appreciated in **Figure 5F** and are consistent with a role for *Bpnt2* in mediating lithium’s effect on GAG sulfation.

## Discussion

In this work, we utilize an *in vitro* fibroblast model system to demonstrate that knockout of a lithium-inhibited enzyme, BPNT2, impairs overall GAG sulfation as well as chondroitin-4-sulfation specifically, and that this impairment stems specifically from a loss of the catalytic activity of BPNT2, as a catalytic-dead construct does not rescue these impairments. The additional missense mutations in BPNT2 that were investigated in this paper are associated with chondrodysplasia in humans and are substitutions of amino acids that are highly conserved across species and happen to occur in close proximity to the catalytic core. We believe that these mutations are therefore likely to interfere with BPNT2 catalysis. Two additional chondrodysplasia-associated BPNT2 mutations have been identified, R187X and S108RfsX48^8,9^. We did not investigate these mutations, but it is understandable that a nonsense mutation and a frameshift mutation would result in more significant disruptions to the protein structure and function. We were interested in how 2 missense mutations, which are less structurally disruptive, are still severe enough to produce a chondrodysplastic disorder. Vissers et al., who initially described these mutations, posited that the mutation of threonine 183 to proline in human *BPNT2* would produce a helix-breaker effect, which would alter secondary BPNT2 structure^8^. They also suggested that the loss of the charged aspartic acid side chain in the D177N mutation would affect the binding of metal cations at the active site^8^. In this work, we have presented data recapitulating impairments in chondrogenesis caused by these two missense mutations, and have further identified that the D(177/175)N mutation generates an N-glycosylation consensus sequence in both human and mouse *Bpnt2*. We can observe this additional glycosylation in BPNT2^D175N^ MEFs. N-glycosylation is a protein modification that occurs co-translationally^21^, and could therefore affect protein folding. Additional kinetic and protein biochemical assays will be important in understanding precisely how these mutations influence BPNT2 catalysis.

As previously reported, we identified decreases in chondroitin-4-sulfate (4S) and increases in unsulfated chondroitin (0S) as a result of the loss of BPNT2. Because of our finding that mutations near the catalytic site do not rescue these alterations, we attribute them to disrupted BPNT2 catalytic activity. Interestingly, we also identified *increases* in chondroitin-6-sulfate (6S), which were not previously identified in tissues (whole embryo preparations) from *Bpnt2-*KO mice. Alterations in 6S could have gone undetected as 4S is the major species of chondroitin present in MEFs (and in whole-embryo mouse preparations)^1^, while 6S is much less abundant and alterations would be less apparent. Contrasting roles for 4S and 6S have been described, wherein 4S is more abundant in developing cartilage, while 6S is more abundant in mature and articular cartilage^22^. In the nervous system, 6S is more abundant in the developing brain, where it is important for synaptic plasticity, whereas 4S is more abundant in the adult brain, where it is thought to promote synaptic stability^23,24^. Importantly, decreases in 6S have been identified in the brains of human patients with bipolar disorder^17^ and lithium treatment is associated with increases in 6S^17^, thought to be corrective. We now present evidence that inhibiting the catalytic activity of a molecular target of lithium can increase 6S levels in a mammalian cell model. The increase in 6S could be a result of shunting or flux given the reductions in 4S and significant elevation of 0S. It is notable that GAG sulfation overall was still decreased in response to the loss of BPNT2 activity, as measured by DMMB assay, which is not specific for any species of sulfated GAG.

While our results clearly indicate that BPNT2 activity is required for altering sulfated GAG/chondroitin sulfation, we are not able to distinguish exactly how this is accomplished. Is it through altering the production of 5-AMP or through defects in the consumption of PAP? In our previous studies of BPNT2’s cytosolic counterpart BPNT1, we found clear evidence of PAP accumulation in mutant animals and cells, which resulted in metabolic toxicity^4,5,19^. An accumulation of PAP could also feedback-inhibit sulfotransferases^25^, impairing GAG sulfation. However, we did not observe an increase of PAP in *Bpnt2*-KO cells. A potential explanation for this is that PAP only needs to locally accumulate within the Golgi, where BPNT2 is located, in order to effectively inhibit sulfotransferases. BPNT1 is still metabolizing PAP in the cytosol. The relative proportion of PAP accumulation in the Golgi to the total amount of PAP in the cell may be too small for such a difference to be detectable by our assays. Isolating only the PAP that is located within the Golgi-lumen would be a technically complex undertaking beyond the scope of this work.

If the phenotypes observed in BPNT2 mutants are due to failure to produce 5’-AMP then it is possible that PAPS transport into the Golgi is impaired. Studies of the PAPS transporter, which moves the sulfate donor PAPS from the cytosol into the Golgi, indicate it is a member of the antiporter family of proteins that may exchange PAPS for 5’-AMP^26,27^. The diminished production of 5’-AMP in the Golgi resulting from loss of BPNT2 activity could prevent PAPS from entering the Golgi and being utilized for sulfation reactions. Further work will be required to determine which, if either, of these mechanisms underlies the observed impairments in sulfation.

In this work, we also show that treatment of MEFs with lithium chloride alters sulfation, and that this alteration is dependent on the presence of BPNT2. Prior work suggests that lithium negatively regulates chondroitin sulfate proteoglycans which would otherwise prevent axon regeneration after spinal cord injury^16^. It has also been shown that lithium promotes neurite outgrowth^28,29^, a process which is normally restricted by chondroitin sulfate^30^. Consistent with these prior findings, we identified decreases in sulfated GAG when MEFs were treated with lithium. Because *Bpnt2*-KO MEFs altered sulfation of GAGs in a manner that phenotypically mimicked WT MEFs treated with lithium, and did not exhibit additive decreases in intracellular GAG sulfation when treated with lithium, we propose that BPNT2 mediates lithium-induced reduction in GAG sulfation. When specifically analyzing chondroitin disaccharides, we identified alterations in WT MEFs treated with lithium consistent with the effects previously identified in *Bpnt2-*KO MEFs, but we did not identify additional alterations when *Bpnt2-*KO MEFs were treated with lithium. These data are consistent with the hypothesis that effects of lithium on chondroitin sulfation are mediated by BPNT2.

The field of psychopharmacology has yet to reach a consensus on how lithium remains so effective in treating bipolar disorder, despite a vast array of other recent psychopharmacologic advances. While this work does not decisively establish lithium’s mechanism of action, it does present evidence that loss of the catalytic activity of known target of lithium, BPNT2, mediates decreases in sulfated GAG, and observable lithium-mediated decreases in sulfation may be due to the loss of BPNT2. On the whole, this work provides a basis for an interesting new hypothesis into how lithium could elicit its psychiatric effects.

## Experimental Procedures

### MEF harvesting and culture methods

Cells were obtained from the *Bpnt2-*knockout mouse line previously generated by our laboratory and available through Jackson Laboratory mouse repository (Jackson #012922). Details regarding the development of this mouse line can be found in Frederick et al. 2008^1^. Mouse embryonic fibroblasts were obtained from E12.5 pups from heterozygous (*Bpnt2*+/−) breeding pairs. Cells from each pup were genotyped (according to Frederick et al. 2008) and cultured in DMEM +4.5 mg/dl glucose and L-glutamine (Gibco), with 10% FBS and 1% penicillin/streptomycin (basal medium). Cells were incubated at 37C with 5% CO2 for the duration of culture. Cells were immortalized by lentiviral expression of SV40 large and small T antigens—briefly, SV40 T antigen lentiviral plasmid (Addgene #22298) was packaged in HEK 293T cells using helper plasmids pMD2.G (Addgene #12259) and psPAX2 (Addgene #12260). 48 hours after transfection of all three plasmids, 3 mL of cell media was collected, filtered through a 0.45 um sterile filter, and added directly to MEF culture medium containing 8 ug/mL polybrene.

### Promotion of chondrogenesis in MEFs

To enhance production of GAGs while in three-dimensional culture, basal medium was changed to chondrogenic medium: DMEM +4.5 mg/dl glucose and L-glutamine (Gibco), with 10% FBS, 1% penicillin/streptomycin, 1% ITS+ supplement (Corning), 0.1 uM dexamethasone, 200 uM ascorbic acid, and 10 ng/mL TGF-β1 (Peprotech). For chondrogenic three-dimensional pellet culture, approximately 1 million cells were seeded in sterile screw-cap 1.7-mL conical tubes and centrifuged at 500g for 5 minutes to form pellets. Cells were cultured as pellets in these tubes with the caps loosened. Pellet media (700 uL per tube) was changed 2 times weekly until cells were harvested for downstream analysis, 7-14 days after seeding. For lithium experiments, medium contained 10 mM LiCl or NaCl for the duration of pellet culture.

### Generation of mutant Bpnt2 MEF lines

Mouse *Bpnt2* cDNA was cloned into the pBABE-puro (Addgene #1764) retroviral vector using BamHI and SalI restriction sites. The plasmid was subsequently mutagenized using traditional site-directed mutagenesis methods to generate D108A, T181P, and D175N mutants, and mutagenesis was verified by Sanger sequencing. Retroviral vectors were each co-transfected with VSV.G (Addgene #14888) and gag/pol (Addgene #14887) vectors into HEK 293T cells using GenJet DNA transfection reagent (SignaGen). 2 mL of viral supernatant was harvested 48 hours post transfection, filtered to remove cells, and added directly to separate *Bpnt2-*KO MEF cultures containing 8 ug/mL polybrene (Invitrogen). Approximately 24 hours post viral transduction, 3 ug/mL puromycin was added to kill non-transduced cells. Cells were incubated in puromycin media for 3 days before being passaged into media without puromycin. Efficacy of transduction was confirmed by measuring *Bpnt2* mRNA and protein expression.

### Quantitative PCR

RNA was collected from culture samples using Qiagen RNeasy Mini Kit, including treatment with Qiagen DNase I. cDNA was synthesized from 1 ug of RNA using iScript cDNA Synthesis Kit from Bio-Rad. PCR reaction was carried out using SsoAdvanced Universal SYBR Green Supermix (Bio-Rad) according to manufacturer instructions. *Bpnt2* mRNA expression (F: 5’-CGCCGATGATAAGATGACCAG-3’ and R: 5’-GCATCCACATGTTCCTCAGTA-3’) was normalized to HPRT mRNA expression (F: 5’-GCAGTACAGCCCCAAAATGG-3’ and R: 5’-ATCCAACAAAGTCTGGCCTGT-3’) to determine relative transcript enrichment.

### Immunoblotting

Protein was collected from cells lysed in RIPA buffer with protease inhibitor (Roche). Protein extracts were passed through a 25g needle to break up DNA and subsequently quantified using BCA assay. 10 ug of total protein was loaded per lane onto a 12% SDS-PAGE gel (Bio-Rad), which was run at 100 V for 2 hours. Proteins were transferred to a 0.2 um PVDF membrane (Bio-Rad) using TransblotTurbo (Bio-Rad) at 1.3A for 7 minutes. Blots were incubated in primary antibodies (sheep anti-BPNT2, 1:1000, Invitrogen #PA5-47893; mouse anti-Actin, 1:1000, Invitrogen #MA1-744) overnight at 4C, washed 3 times in 0.1% TBS-Tween then in secondary antibodies (AlexaFluor680 anti-sheep 1:20,000; AlexaFluor800 anti-mouse 1:20,000) for 2 hours at room temperature and washed 3 times in 0.1% TBS-T. Blots were imaged on LiCor Odyssey.

### PNGaseF digestion

10 ug of protein extract were digested with PNGaseF (NEB) according to manufacturer instructions. Full digest products were subsequently run on gel as described above.

### Measurement of PAP in MEFs

Briefly, approximately 1 million cell MEF pellets which had been cultured as pellet for 14 days were boiled for 3 min in 150 μL of PAP isolation buffer (50 mM glycine, pH 9.2) and disrupted mechanically with a tissue pestle. Homogenates were clarified by centrifugation at 16100 × g, 4 °C for 20 min. Then 0.2 volumes of chloroform (CHCl_3_) was added, and samples were vortexed to mix. Samples were again centrifuged at 16100 × g, 4 °C for 20 min. The upper aqueous phase was then collected and used for the assay. To quantify PAP levels, we used a colorimetric microplate absorbance assay in which recombinant mouse SULT1A1-GST is used to transfer a sulfate group from p-nitrophenyl sulfate to 2-naphthol, using PAP as a catalytic cofactor^31^. Briefly, 20 μL of the tissue lysate was incubated with 180 μL of PAP reaction mixture [100 mM bis·Tris propane (pH 7.0), 2.5 mM β-mercaptoethanol, 2.5 mM p-nitrophenyl sulfate, 1 mM β-naphthol, and 1 μg of PAP-free recombinant mouse SULT1A1-GST]. Reactions velocities were determined by monitoring the production of 4-nitrophenol at 405 nm. Concentrations of PAP in lysates were determined by comparing reaction rates acquired from kinetic analysis to those of a series of PAP standards run concurrently on the same plate.

### Dimethylmethylene blue (DMMB) assay

Media was collected from cell pellets upon harvest and kept at −20C until used for downstream analyses. After removing media, cell pellets were rinsed in 1X PBS. Pellets were then incubated in 300 ul 10mM Tris-HCl (pH 7.5) solution containing 100ug/mL proteinase K (Roche) at 60C overnight followed by 30 min at 90C to denature Proteinase K. 40 ul of each sample was loaded onto a clear-bottom 96-well plate in duplicate, and 200 ul of pH 1.5 DMMB reagent (prepared according to Zheng and Levenston 2015^32^) was added using a multichannel pipette. Absorption was immediately measured at 525 and 595 nm, and 595 measurement was subtracted from 525 to yield final reading. Quantity of sulfated GAG was determined by comparison to a standard curve of bovine chondroitin-4-sulfate (Sigma) prepared in 10mM Tris-HCl. Amount of sulfated GAG was normalized to cell count across samples. For analysis of secreted sulfated GAG in media, 60ul media was added directly to clear-bottom 96-well plate in duplicate, and readings were compared against bovine chondroitin-4-sulfate standards prepared in culture medium.

### High performance liquid chromatography analysis of chondroitin-sulfate disaccharides

1-million-cell pellets were homogenized in 400 ul GAG preparation buffer (50 mM Tris, pH 8.0, 10 mM NaCl, 3 mM MgCl_2_), containing 4 ul of 2mg/mL Proteinase K, using tissue pestle. Sample was incubated overnight at 56C. After digest, the samples were heated at 90C for 30 minutes to denature Proteinase K. Precipitated material was separated by centrifugation. Sample buffer was changed to 0.1 M ammonium acetate, pH 7.0 by using 3 kDa Millipore concentrator by concentration/dilution until initial concentration of homogenization buffer decreased 500-times. The volume of concentrated samples was adjusted to 70 ul and 3 ul of Chondroitinase ABC (1.4 U/ml Stock solution, containing BSA, Seikagaku) was added to each sample. Reaction mixture was incubated at 37C for 4 hours. After chondroitinase ABC cleavage, 130 ul of water was added. Released disaccharides were filtered through a 10 kDa concentrator. This procedure was repeated once more to improve yield. GAG samples were lyophilized using a SpeedVac at 25 °C overnight. Lyophilized samples were stored at −80C until fluorescent labeling. Labeling of disaccharides was performed with 2-aminobenzamide (2-AB) by published procedure^33^. An aliquot of 5-7 ul of labeling mixture (0.35 M 2-AB, 1 M NaCNBH_3_ solution in 30% acetic acid in dimethyl sulfoxide) was added to lyophilized samples or disaccharide standards and the mixture was incubated for 3 hours at 65C. Labeling reaction mixtures were spotted on a strip of Whatman 3MChr paper and washed with 1 ml of acetonitrile six times. Cleaned disaccharides were eluted with three aliquots of 50, 75 and 75 ul of water by using 0.2 um centrifugal device. The analysis of labeled disaccharides was performed by HPLC. The HPLC system included Waters 515 Pumps, Waters 517plus Autosampler, Waters Pump Control Module II and Shimadzu RF-10Axl spectrofluorometer detector under Waters Empower software. Sample analysis was performed on Supelco-LC-NH_2_ 25 cm × 4.6 mm (Sigma). Column was equilibrated with 16 mM NaH_2_PO_4_ with flow rate of 1 ml/min. The samples of 50-100 ul were injected and eluted with 60 min linear gradient 16 mM – 800 mM NaH_2_PO_4_ with flow rate of 1 ml/min as in Yoshida et al. 1989^34^. Disaccharide elution was monitored by fluorescence at 420 nm with excitation at 330 nm. Peaks were identified by comparison to 4S, 6S, and 0S chondroitin standards (Sigma). Calculations were determined by integrating each peak on the resultant chromatogram and calculating ratios of chondroitin species. Chromatograms were analyzed by Empower software.

## Data availability

All relevant data are contained within the manuscript. Additional information may be requested by contacting the corresponding author.

## Acknowledgements

The authors would like to thank Sun Peck, Bradley Clarke, Garrett Kaas, Lucia Plant, and Jane Wright for their insights and assistance throughout the course of this work.

## Author Contributions

B.S.E and J.D.Y. conceived of the experiments and wrote the manuscript. B.S.E. designed and performed the majority of the experiments described herein, Z.A.L. performed HPLC analyses, A.J.W, F.Y., and A.T.H. helped to carry out cell culture experiments.

